# The role of subicular VIP-expressing interneurons on seizure dynamics in the intrahippocampal kainic acid model of temporal lobe epilepsy

**DOI:** 10.1101/2023.05.30.542857

**Authors:** Sadegh Rahimi, Pariya Salami, Pawel Matulewicz, Armin Schmuck, Anneliese Bukovac, Arnau Ramos-Prats, Ramon Osman Tasan, Meinrad Drexel

**Author notes:** Corresponding author, Priv.-Doz. Mag. Meinrad Drexel, PhD, Medical University of Innsbruck, Institute of Pharmacology, Peter-Mayr-Str. 1/1a, 6020 Innsbruck, Austria, Phone: +43(0)512/9003-70405. Declarations of interest: none.

## Abstract

The subiculum, a key output region of the hippocampus, is increasingly recognized as playing a crucial role in seizure initiation and spread. The subiculum consists of glutamatergic pyramidal cells, which show alterations in intrinsic excitability in the course of epilepsy, and multiple types of GABAergic interneurons, which exhibit varying characteristics in epilepsy. In this study, we aimed to assess the role of the vasoactive intestinal peptide interneurons (VIP-INs) of the ventral subiculum in the pathophysiology of temporal lobe epilepsy. We observed that an anatomically restricted inhibition of VIP-INs of the ventral subiculum was sufficient to reduce seizures in the intrahippocampal kainic acid model of epilepsy, changing the circadian rhythm of seizures, emphasizing the critical role of this small cell population in modulating TLE. As we expected, permanent unilateral or bilateral silencing of VIP-INs of the ventral subiculum in non-epileptic animals did not induce seizures or epileptiform activity. Interestingly, transient activation of VIP-INs of the ventral subiculum was enough to increase the frequency of seizures in the acute seizure model. Our results offer new perspectives on the crucial involvement of VIP-INs of the ventral subiculum in the pathophysiology of TLE. Given the observed predominant disinhibitory role of the VIP-INs input in subicular microcircuits, modifications of this input could be considered in the development of therapeutic strategies to improve seizure control.

## Introduction

Accumulating evidence from both clinical and experimental studies has provided insights into the molecular changes, hypersynchrony, and abnormal anatomical structure of the subiculum that contribute to epileptic seizures (1). Over two decades ago, in brain slices obtained from patients with temporal lobe epilepsy (TLE), a spontaneous rhythmic activity exclusively originating in the subiculum was observed. These synchronous events resembled the interictal discharges recorded in the patients’ electroencephalograms (EEG) (2). Analysis of resected hippocampi from individuals with TLE revealed no significant differences in the spontaneous epileptiform activities in the subiculum between sclerotic and non-sclerotic tissues (3). This suggested that modifications in synaptic transmission and intrinsic properties of the subiculum play a crucial role in the development of epilepsy, rather than deafferentation-induced alterations. Consistent with clinical findings indicating relatively preserved integrity of the subiculum in TLE (4), animal models of TLE also demonstrated less noticeable neuron loss in the subiculum compared to the CA1/ CA3 regions (5).

The subiculum consists of glutamatergic pyramidal cells (PCs) that undergo alterations in intrinsic excitability during the course of epilepsy, and various types of GABAergic interneurons (GABAergic-INs) that exhibit diverse characteristics in epilepsy (6–8). Although GABAergic-INs constitute only 10% to 15% of the total neuronal population in the hippocampus (9), their extensive anatomical and physiological diversity enables them to exert significant regulatory influence on cellular and circuit functions (10). The dysfunction of subicular INs in epilepsy can be simplified as an imbalance of excitation and inhibition (E/I). However, these INs form intricate circuit patterns, including feedforward, feedback, and disinhibitory connectivity (11). Consequently, the three major subclasses of subicular INs - parvalbumin-expressing (PV-INs), somatostatin-expressing (SOM-INs), and vasocactive intestinal polypeptide-expressing (VIP-INs) - may play opposing roles in the pathophysiology of TLE. Animal models of TLE have demonstrated selective degeneration of subicular and CA1 PV-INs (12, 13). Moreover, silencing PV-containing basket cells and axo-axonic cells in the subiculum led to recurrent series of interictal spikes and spontaneous seizures in over 60 percent of the animals (14). Loss of SOM-INs in the CA1 region of the hippocampus following pilocarpine-induced seizures has also been reported (15). Additionally, decreased dendritic inhibition of PCs resulting from permanent functional silencing of subicular SOM-INs caused spontaneous seizures in approximately one-third of the animals (16). While the role, changes in the numbers, and properties of subicular PV-INs and SOM-INs in epilepsy are well documented (1), the impact of subicular VIP-INs in epilepsy remains poorly understood.

Hippocampal VIP-INs can be classified into two main groups based on their targets (17). The first group consists of VIP-basket cells (VIP-BCs), which co-express cholecystokinin (CCK) and primarily inhibit the somatic region of PCs. The second group comprises interneuron selective interneurons (IS INs), which selectively target other INs. IS INs can be further divided into two subtypes: (a) Interneurons co-expressing calretinin (CR), with cell bodies located in the stratum pyramidale (SP) or near it, projecting to the stratum oriens/alveus border (referred to as VIP-IS O/A INs or IS type III). These interneurons mainly target SOM-expressing oriens lacunosum-moleculare (OLM) interneurons. (b) VIP-INs that project their axons to the stratum radiatum (SR), with cell bodies located either at the SR/ stratum lacunosum-moleculare border (IS type II) or at SR/SP. These VIP-INs target interneurons that mainly innervate proximal dendrites of PCs in stratum radiatum (17–19).

In human TLE, the loss of pyramidal neurons has been associated with upregulation of VIP receptors in the hippocampus, while the distribution and pattern of VIP-INs remained unchanged (20). Children with chronic epilepsy have been found to exhibit elevated levels of VIP in their cerebrospinal fluid (21). Moreover, in focal cortical dysplasia (type IIb), primary common cause of medically refractory pediatric epilepsy, there is a notable increase in the ratio of VIP-INs to other interneurons (22). In rodent models, increased firing rates of VIP-INs were observed in cortical slices treated with tubocurarine as spikes and waves developed (23). Interestingly, most VIP-INs are preserved in animal models of epilepsy (24, 25), although the OLM interneurons targeted by VIP-INs are selectively lost (26).

While there is growing evidence suggesting the potential significance of VIP-INs in TLE, there is a lack of consensus regarding their role. The upregulation VIP receptors or lack of change in VIP-INs can be interpreted as either causative (22, 24) or compensatory (27) in the context of epileptic excitotoxicity. Additionally, no studies have explored the functional role of VIP-INs specifically in the subiculum. Although VIP-INs are relatively sparse compared to PV-INs and SOM-INs, their unique connectivity patterns suggest they may play a key role in the subicular circuit. However, their precise function in epilepsy remains unclear.

To investigate the potential role of VIP-INs of the ventral subiculum in the development and maintenance of TLE, we employed a targeted approach to inhibit neurotransmitter release from these neurons. Our study involved both healthy control mice and a chronic mouse model of epilepsy, with continuous EEG monitoring using telemetric recording. In mice with epilepsy, the silencing of VIP-INs of the ventral subiculum leads to a reduction in the severity of seizures, while silencing them in healthy mice does not induce epileptiform activity. Furthermore, we conducted experiments using chemogenetic activation of these interneurons in a model of acute seizures, which resulted in an increase in epileptiform activity. These findings provide novel insights into the significant involvement of VIP-INs of the ventral subiculum in the pathophysiology of TLE.

## Results

### Silencing of VIP-INs of the ventral subiculum in epileptic mice dampens seizures

Consistent with the CA1 region (17), we also identified three distinct subgroups of VIP-INs in the ventral subiculum (*n* = 3, VIP:Ai9-tdTomato mice). These subgroups include INs expressing VIP alone (IS type II), INs co-expressing VIP and CCK (VIP-BCs), and INs co-expressing VIP and CR (IS type III; Figure 1A). Next, we conducted experiments to examine whether dysfunction of VIP-INs of the ventral subiculum can influence seizure severity and epileptogenesis. For this purpose, we utilized an animal model of TLE induced by intrahippocampal injection of kainic acid (IH-KA, 200 ng). Telemetric EEG transmitters were surgically implanted in 16 VIP-cre mice (8 males and 8 females). The animals were randomly assigned to two groups, with one group receiving AAV-GFP and the other group receiving AAV-TeLC, after a 7-day period (Figure 1B). Prior to vector injection, the severity of status epilepticus (SE) and subsequent spontaneous recurrent seizures were evaluated to establish the baseline epileptic activity for both experimental groups. During SE on Day 1, the median (range) number of seizures observed was 10 (1 to 19) and 8.5 (1 to 14) for the GFP and TeLC groups, respectively (Figure 1D). No significant difference was observed between the groups based on the number of seizures during SE. Since seizures during SE may not always exhibit postictal depression (Supplementary Figure 1B), the number of seizures could be underestimated. Therefore, in addition to seizures, we also compared the groups based on the number of epileptiform spikes and spike trains. During SE, no significant differences were observed between the groups regarding the number of epileptiform spikes (Supplementary Figure 1C-D) or spike trains (Supplementary Figure 1E-F). Additionally, during the six days of recording after SE and before viral vector injection, the number of spontaneous recurrent seizures (SRSs) was similar between the two experimental groups (Figure 1E). Following a two-week waiting period after viral vector injection, the number and duration of seizures were assessed and compared between the groups, revealing a significant difference (Figure 1F). Taking into account the inter-individual differences in the number of seizures in the IH-KA model, we further evaluated the seizure ratio for each animal by comparing the number of seizures before and after vector injection. The median (range) seizure ratio was 0.55 (−1.00 to 0.86) for the GFP group and −0.33 (−1.00 to 0.42) for the TeLC group (Figure 1G). GFP-injected mice generally exhibited an increasing trend in the number of seizures, whereas the TeLC group showed a decreasing ratio, indicating a significant difference (*p* < 0.05) between the two groups.

**Figure 1.**
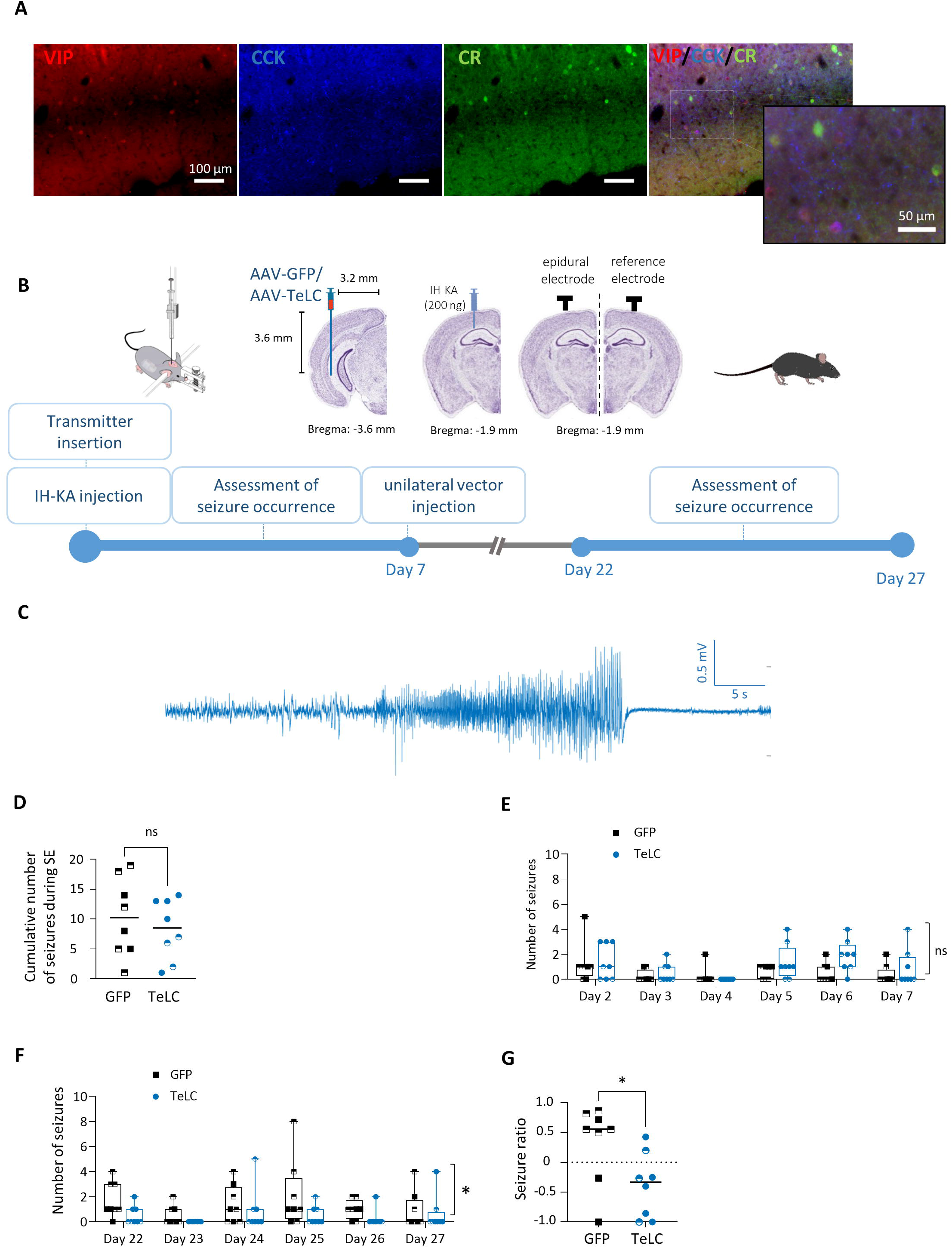
Silencing VIP-INs of the ventral subiculum in epileptic mice reduce the number of seizures. (A) Representative example of triple labeling in brain slices from the ventral subiculum of VIP:Ai9-tdTomato mice (*n* = 3). (B) Study protocol and injection coordinates for intrahippocampal KA (IH-KA, 200 ng), AAV-TeLC-GFP (*n*=8), and AAV-GFP (*n*=8). Seizure frequency and duration were evaluated on Day one (SE), the six days following KA injection (Day 2 to Day 7), as well as on the six days commencing two weeks after vector injection (Day 22 to Day 27). (C) Representative seizure recorded 3 days after IH-KA injection. Seizures are typically preceded by a series of epileptiform spikes and were identified as continuous activity lasting at least 10 seconds with an amplitude at least twice that of the baseline, accompanied by postictal depression. (D) During SE, the median (range) number of seizures was 10 (1 to 19) for the GFP group (*n*=8) and 8.5 (1 to 14) for the TeLC group (*n*=8), with no significant difference between the groups (two-tailed Mann-Whitney test; *p* = 0.625). (E) The number of seizures during Day 2 to Day 7 (before vector injection). Both groups developed spontaneous seizures. A two-way ANOVA test revealed a significant difference between days (*F* = 4.879, *p* = 0.005), but not between groups (*F* = 1.829, *p* = 0.198, *n* = 8 for both GFP and TeLC groups). (F) The number of seizures during Day 22 to Day 27 (after vector injection). Both groups had a stable daily number of seizures (*F* = 1.540, *p* = 0.188, no significant difference between days, according to a two-way ANOVA test). However, the TeLC group had a significantly lower number of seizures compared to the GFP group (*F* = 6.296, *p* = 0.0250, *n* = 8 for both groups). (G) The median (range) seizure ratio was 0.55 (−1.00 to 0.86) for the GFP group (*n*=8) and −0.33 (−1.00 to 0.42) for the TeLC group (*n*=8), with a significant difference according to the two-tailed Mann-Whitney test (*p* = 0.022). Male individuals are represented by half-filled shapes in both groups. Mouse coronal slices in (B) are adapted from the Allen Mouse Brain Atlas, available online: http://atlas.brain-map.org/atlas. The walking mouse (right) is sourced from https://scidraw.io.

We next evaluated the time spent in seizure in both groups. The median (range) cumulative duration of seizures during SE was 314.1 s (10.34 s to 1098 s) for the GFP group and 266.10 s (17 s to 703 s) for the TeLC group (Figure 2A), showing no significant difference between the groups. Before vector injection, the time spent in seizures was comparable between the experimental groups (Figure 2B). However, following vector injection, a significant difference in seizure duration was observed between the groups (*p* < 0.05, Figure 2C). Additionally, we evaluated the seizure duration ratio for each animal, revealing a median (range) of 0.66 (−1.0 to 0.96) for GFP mice and −0.36 (−1.0 to 0.34) for TeLC mice, indicating a significant difference (*p* < 0.05, Figure 2D).

**Figure 2.**
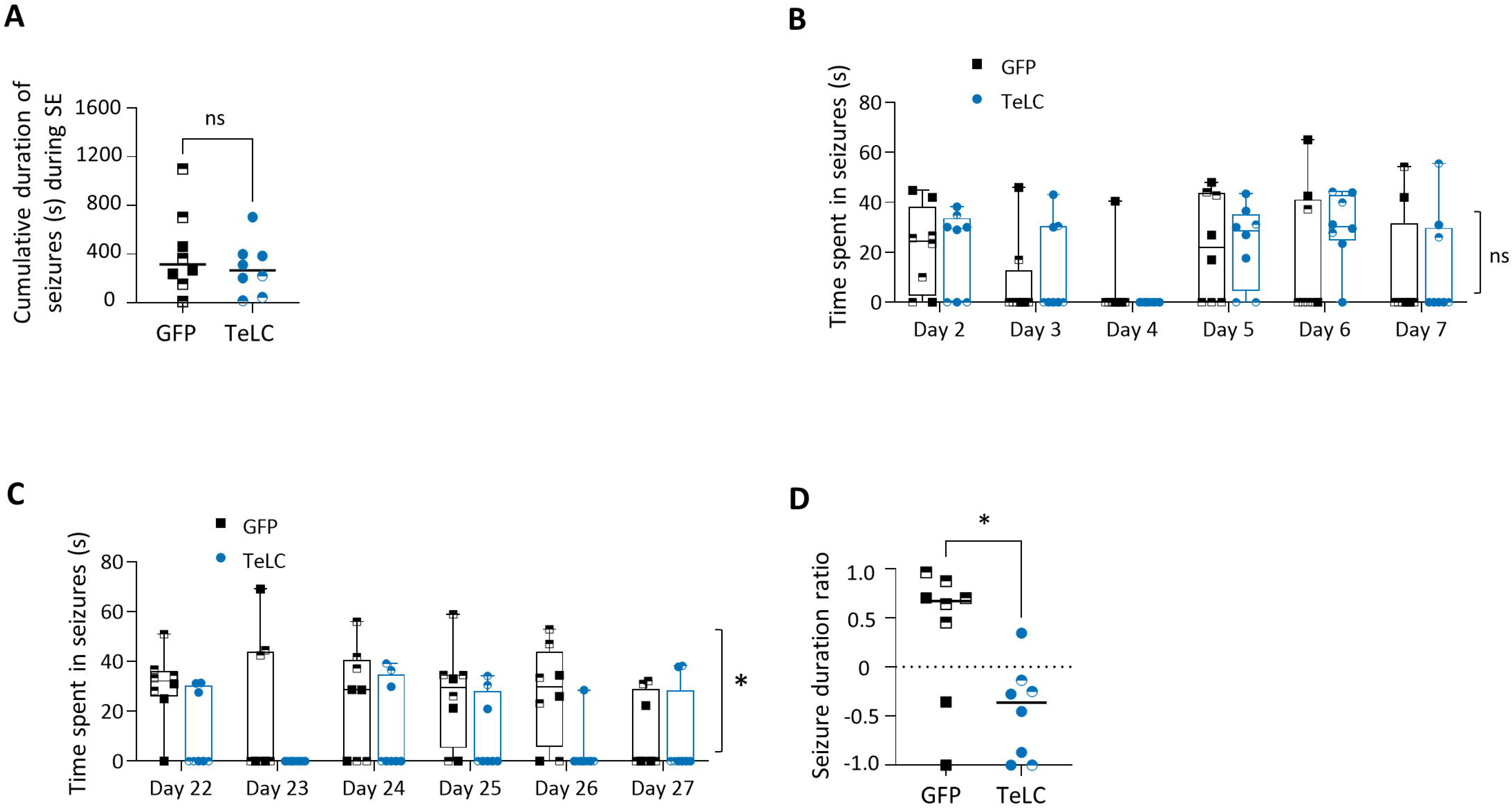
Silencing VIP-INs of the ventral subiculum in epileptic mice reduce the duration of seizures. (A) During SE, the median (range) cumulative duration of seizures was 314.1 s (10.34 s to 1098 s) for the GFP group (*n*=8) and 266.10 s (17 s to 703 s) for the TeLC group (*n*=8), with no significant difference between the groups according to the two-tailed Mann-Whitney test (*p* = 0.645). (B) The daily time spent in seizures during Day 2 to Day 7 (before vector injection). Both groups developed spontaneous seizures. There was a significant difference in the time spent in seizures between days (*F* = 4.561, *p* = 0.005), but no significant difference between the GFP and TeLC groups (*F* = 1.695, *p* = 0.687, *n* = 8 for both groups). (C) The daily time spent in seizures during Day 22 to Day 27 (after vector injection). Both groups maintained stable values for the time spent in seizures (*F* = 1.286, *p* = 0.287, no significant difference between days, according to two-way ANOVA test); however, the mice in the TeLC group spent significantly less time seizures compared to the GFP group (*F* = 10.350, *p* = 0.006, *n* = 8 for both groups). (D) The median (range) seizure duration ratio was 0.66 (−1.0 to 0.96) for GFP mice (*n*=8) and −0.36 (−1.0 to 0.34) for TeLC mice (*n*=8), with a significant difference according to the two-tailed Mann-Whitney test (*p* = 0.027). Male individuals are represented by half-filled shapes in both groups.

Upon evaluating circadian changes in the seizure ratio for both groups (Figure 3B), a significant reduction was observed during the light phase (*p* < 0.05). However, this reduction was not significant during the dark phase (Figure 3C). Notably, the reduction in time spent in seizures was more pronounced during the light phase (*p* < 0.05, Figure 3D), aligning with the trend observed for the number of seizures. The vigilance state of the animals was assessed both before and after vector injection. Interestingly, it was found that the alteration in the circadian rhythm of the seizures was not correlated with changes in the vigilance state. Furthermore, no significant changes in sleep macro and micro architecture were observed that corresponded to these seizure rhythm changes (Supplementary Figure 2).

**Figure 3.**
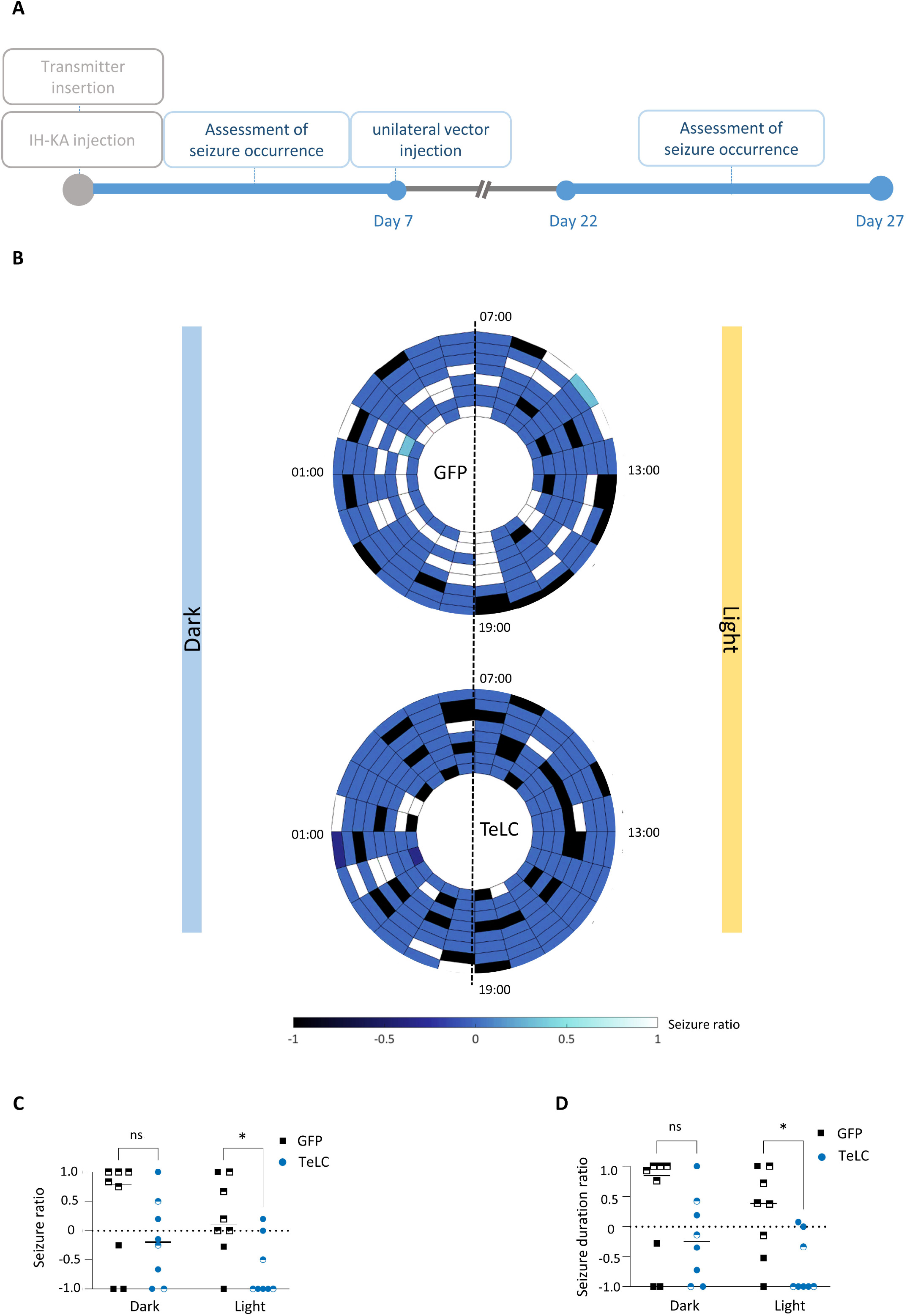
More prominent seizure reduction during the light phase. (A) The seizure ratio was calculated for each animal by comparing the number of seizures after vector injection (Day 22 to Day 27) to the number of seizures before vector injection (Day 2 to Day 7). (B) Each cycle represents a single animal from either the GFP (*n* = 8) or TeLC group (*n* = 8), with each rectangle indicating the seizure ratio during one hour of the 24-hour day. (C) The reduction in the number of seizures was more prominent during the light phase (*p* = 0.044, post-hoc Šídák test) compared to the dark phase (*p* = 0.369, post-hoc Šídák test) according to two-way ANOVA test. (D) The decrease in the cumulative duration of seizures was more pronounced during the light phase (*p* = 0.042, post-hoc Šídák test) compared to the dark phase (*p* = 0.323, post-hoc Šídák test) according to the two-way ANOVA test. Male individuals are represented by half-filled shapes in both groups.

The outcomes of the first experiment indicated that silencing VIP-INs in the ventral subiculum effectively reduced both the frequency and duration of seizures, particularly during the light phase.

### Silencing VIP-INs of the ventral subiculum in healthy mice does not induce seizures or epileptiform activity

In our previous research, we reported that silencing PV-INs (14) and SOM-INs (16) of the ventral subiculum results in the development of spontaneous recurrent seizures (SRSs). However, our current findings demonstrate that silencing VIP-INs of the ventral subiculum can actually alleviate seizures in epileptic animals. Therefore, based on these findings, if we were to silence VIP-INs of the ventral subiculum in healthy mice, it is unlikely to induce any SRSs. To investigate this, we analyzed EEG patterns in healthy male VIP-cre mice unilaterally injected with either AAV-TeLC (*n* = 5) or a control AAV-GFP (*n* = 5) viral vector into the ventral subicular (Figure 4A). Telemetric EEG recordings were conducted continuously for 50 days, and the recorded EEG traces were visually reviewed to identify seizures (see Methods). No seizures were detected in any of the AAV-TeLC- or AAV-GFP-injected animals (Figure 4B). At the injection site, the median (range) percentage of the VIP-INs expressed GFP was 46.67% (41.18% to 90%), while 95.84% (82.35% to 100%) of the GFP-expressing INs also co-expressed VIP (Figure 4C), highlighting the specificity of the experimental manipulation using the TeLC vector.

**Figure 4.**
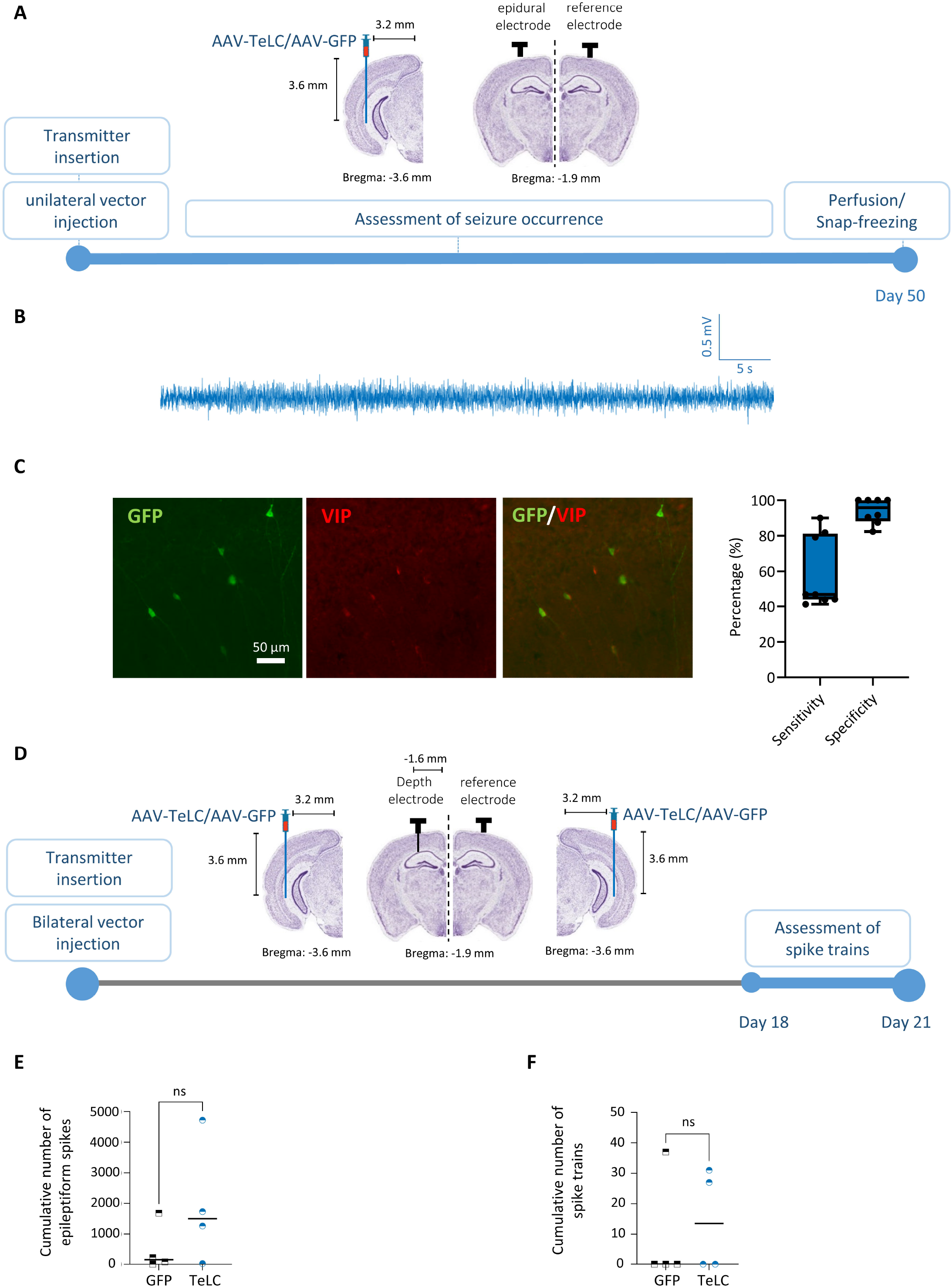
Silencing of VIP-INs of the ventral subiculum does not induce seizures in healthy mice. (A) Study protocol and injection coordinates for intrahippocampal AAV-TeLC-GFP (*n*=5) and AAV-GFP (*n*=5) injections in non-epileptic mice, followed by long-term telemetric EEG recordings. (B) Representative 1-minute EEG-trace obtained 14 days after AAV-TeLC injection. No seizures were observed during the 50-day recording period. (C) Rate of GFP expression in cells. At the injection site, the median (range) percentage of VIP-INs expressed GFP (101 cells out of 169 cells) was 46.67% (41.18% to 90%), while 95.84% (82.35% to 100%) of the GFP-expressing interneurons also co-expressed VIP (101 cells out of 108 cells; *n*=4 mice from the GFP group, *n*=4 mice from the TeLC group). (D) Study protocol and injection coordinates for bilateral intrahippocampal AAV-TeLC-GFP (*n*=4) and AAV-GFP (*n*=4) injections in non-epileptic mice, followed by long-term LFP recording from dorsal CA1. No seizures were observed during the 21-day recording period. (E) Median (range) cumulative number of epileptiform spikes (for the entire 24-hour assessment) was 148 (3 to 1687) for the GFP group (*n*=4) and 1500 (24 to 4723) for the TeLC group (*n*=4). There was no significant difference between the groups according to the two-tailed Mann-Whitney test (*p*=0.343). (F) Median (range) number of spike trains was 0.0 (0.0 to 37) for the GFP group (*n*=4) and 13.50 (0.0 to 31) for the TeLC group (*n*=4), with no significant difference between the groups according to the two-tailed Mann-Whitney test (*p* > 0.99). Mouse coronal slices in (A) and (D) are adapted from the Allen Mouse Brain Atlas, available online: http://atlas.brain-map.org/atlas.

To ensure the detection of subtle focal epileptiform activity, continuous local field potential (LFP) recording was conducted in mice that underwent bilateral injection of AAV-TeLC (*n* = 4) or AAV-GFP (*n* = 4) into the ventral subiculum (Figure 4D). The recordings were performed for 21 days, and data were visually reviewed. No seizures were detected in either group. Furthermore, epileptiform spikes and spike trains were assessed using a custom-made MATLAB code for three consecutive days, with 6 hours of recording per day (from 18:00 to 24:00, starting from Day 18). We observed a minimal number of epileptiform spikes and spike trains. The median (range) cumulative number of epileptiform spikes was 148 (3 to 1687) for GFP animals and 1500 (24 to 4723) for TeLC animals (Figure 4E). No significant difference was found between the groups. Additionally, the median (range) number of spike trains was 0.0 (0.0 to 37) and 13.50 (0.0 to 31) for GFP and TeLC animals, respectively (Figure 4F), and there was no significant difference between the groups.

### Activating VIP-INs of the ventral subiculum in epileptic mice increases seizures

If silencing VIP-INs of the ventral subiculum reduces seizures in the chronic model of epilepsy, it is expected that activation of these INs would increase seizure activity. To test this hypothesis, we utilized an experimental model of acute focal seizures induced by IH-KA injection (7 ng). AAV-hM3D(Gq) was injected into 15 VIP-Cre mice (10 males, 5 females), and the acute seizure model was induced on day 10 by IH-KA injection (Figure 5A). The animals were randomly divided into two groups, CNO (*n* = 9) or NaCl (*n* = 6), and received an i.p. injection of CNO or NaCl, respectively, half an hour before the IH-KA injection. All animals developed seizures within the first hour after IH-KA injection (Figure 5B-C). The median (range) number of seizures in the NaCl group was 2 (1 to 4), whereas CNO group exhibited a higher median value of 5 (2 to 7) seizures (Figure 5D). The CNO-injected mice demonstrated a significant higher number of seizures compared to the NaCl group (*p* < 0.05). The median (range) cumulative duration of seizures for the NaCl group was 76.02 s (33.08 s to 150.50 s), while for the CNO group it was 138.70 s (51.89 s to 225.0 s), indicating a trend towards increased seizure duration in the CNO group compared to control mice (Figure 5E). The findings from the acute seizure model suggest that the enhanced activity of VIP-INs during seizure induction is associated with heightened seizure severity.

**Figure 5.**
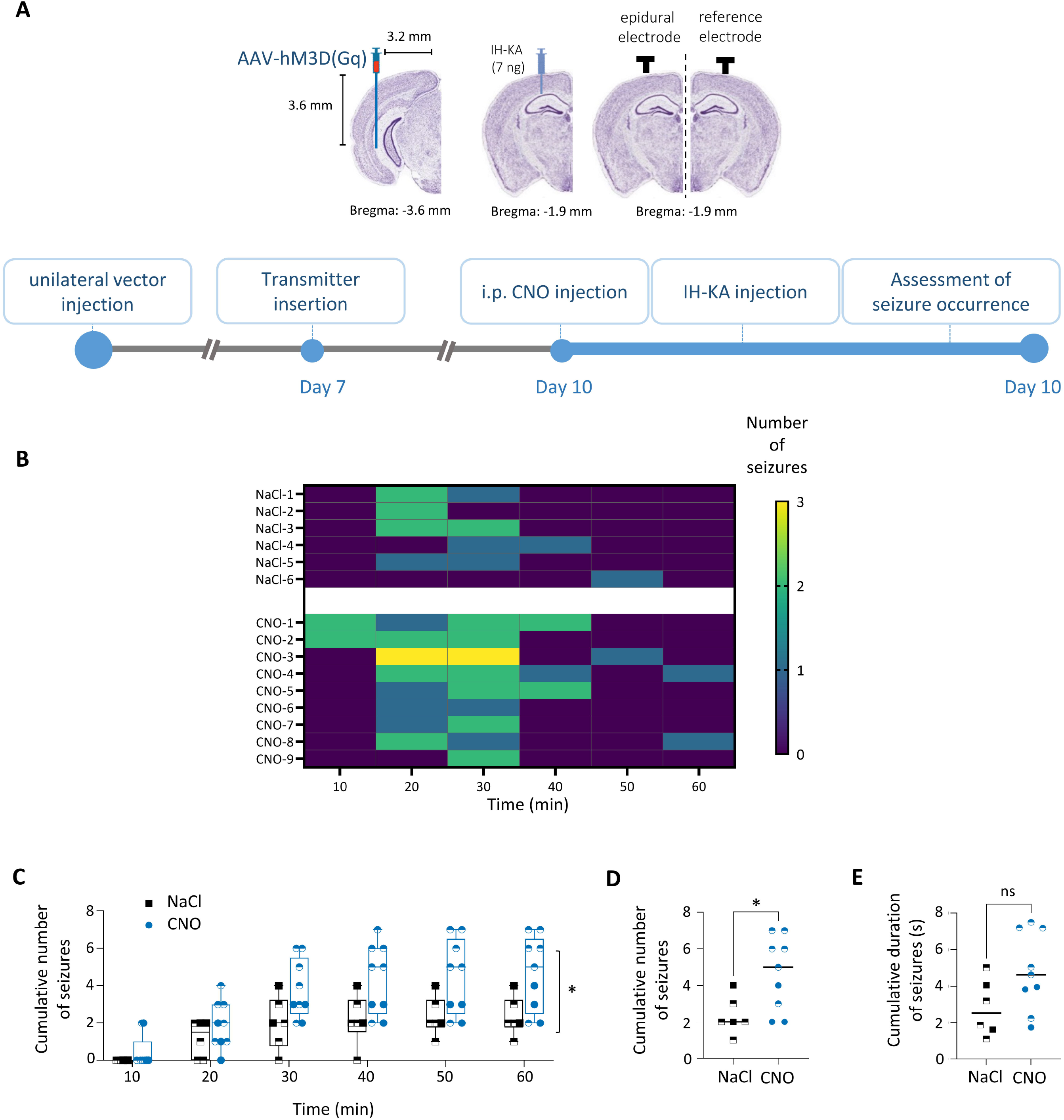
Activating VIP-INs of the ventral subiculum in model of acute seizures results in increased seizure frequency. (A) Study protocol and injection coordinates for AAV-hM3D(Gq) (*n*=15) and intrahippocampal KA (IH-KA, 7 ng) injections. The animals were blindly divided into two groups, NaCl (*n*=6) and CNO (*n*=9), receiving intraperitoneal (i.p.) injection of NaCl and CNO at Day 10, respectively. The i.p. injection was administered 30 minutes prior to the IH-KA injection. (B) Number of seizures for each individual animal in the first 60 minutes after IH-KA injection (10-minute bin). (C) After IH-KA injection, animals in the NaCl (*n*=6) and CNO (*n*=9) groups developed seizures. A significant difference was found between the groups (Two-way ANOVA; *F* = 5.353, *p* = 0.037). (D) Median (range) number of seizures in the NaCl group was 2 (1 to 4), while it was 5 (2 to 7) for the CNO group. A significant difference was reported according to the two-tailed Mann-Whitney test (*p* = 0.034). (E) Median (range) cumulative duration of seizures for the NaCl group was 76.02 s (33.08 s to 150.50 s), and for the CNO group it was 138.70 s (51.89 s to 225.0 s). There were no significant differences between the groups, despite a strong trend (*p* = 0.087, according to the two-tailed Mann-Whitney test). Male individuals are represented by half-filled shapes in both groups. Mouse coronal slices in (B) are adapted from the Allen Mouse Brain Atlas, available online: http://atlas.brain-map.org/atlas.

## Discussion

Epileptic seizures are a result of excessive electrical discharges (28), caused by an imbalance between inhibition and excitation in the brain (29). The subiculum, which serves as a key output region of the hippocampus, is increasingly recognized for its crucial role in seizure initiation and propagation (1). The concept of dynamic excitation/inhibition balance, modulated by circuit disinhibition, has emerged as an important mechanism for information-processing (19). However, the specific role of a distinct subpopulation of subicular GABAergic cells expressing VIP in epilepsy-related disinhibitory modulation has not been investigated in previous studies. Despite the heterogeneity of subicular VIP-INs and their lower numbers compared to PV-INs and SOM-INs (14, 16), selectively inhibiting VIP-INs of the ventral subiculum within a restricted anatomical region proved sufficient to reduce seizures in the chronic model of epilepsy, underscoring the critical role of this small cell population in modulating TLE. As anticipated, permanent unilateral or bilateral silencing of VIP-INs of the ventral subiculum in non-epileptic animals did not induce seizures or epileptiform activity, whereas transient activation of VIP-INs of the ventral subiculum was found to increase the frequency of seizures in the acute seizure model. Our findings offer new insights into the pivotal role of VIP-INs of the ventral subiculum in regulating the excitability of the subicular network within the context of epilepsy. Hippocampal VIP-INs consist of two main subtypes, VIP-BCs and ISs (IS type II and III). These subtypes have opposing roles in hippocampal circuits, with VIP-BCs directly inhibiting PCs while ISs disinhibit PCs (19). Our findings in both healthy and epileptic animals suggest a more prominent role for ISs. In our previous studies, we demonstrated that silencing PV-INs (14) and SOM-INs (16) of the ventral subiculum alone is sufficient of induce seizures. However, this is not the case for VIP-INs of the ventral subiculum, as silencing these neurons in healthy animals did not lead to seizures. This indicates a potential disinhibitory function VIP-INs of the ventral subiculum in the context of epilepsy. Furthermore, silencing of VIP-INs of the ventral subiculum in epileptic animals resulted in a reduction of seizure activity, while activating VIP-INs of the ventral subiculum in an acute model of focal seizures increased the occurrence of seizures. These findings collectively highlight the significant disinhibitory role played by VIP-INs of the ventral subiculum in the context of epilepsy.

To the best of our knowledge, our study represents the first attempt to investigate the involvement of VIP-INs of the ventral subiculum in the development of TLE. However, a limited number of studies have examined both VIP-BCs and ISs in hippocampal sector CA1 in various animal models of epilepsy. In the IH-KA model, it has been reported that the number of CCK-expressing INs (of which VIP-BCs constitute approximately 27% (9)) was reduced in the ventral hippocampus following KA injection, although the surviving CCK-BCs did not exhibit significant changes in their intrinsic and synaptic properties or connectivity with CA1-PCs (30). In the pilocarpine model of epilepsy in mice, the overall density of IS type III INs remained relatively preserved (26, 25), but the number of axonal boutons produced by these cells within the CA1 oriens/alveus was significantly lower after SE (26). Furthermore, VIP+/CR-INs, which include both IS type II and VIP-BCs, decreased to 54% compared to non-epileptic animals (25). However, no study has specifically evaluated the vulnerability of IS type II cells in the context of epilepsy. It may be argued, that VIP-BCs are more susceptible to damage than ISs in the IH-KA model of epilepsy. Consequently, the balance of cell types may shift, leading to a greater impact from ISs when VIP-INs are silenced in the chronic model of epilepsy. However, the results of silencing VIP-INs in non-epileptic animals and in the model of acute seizures support the predominant role of ISs, regardless of the number of viable cells.

The role of VIP-INs of the ventral subiculum in TLE can be influenced not only by the release of GABA from these INs, but also by endogenous VIP. In the hippocampus, VIP is exclusively expressed by GABAergic-INs (31) and interacts with two receptors, VPAC1 and VPAC2, as well as PAC1 with lower affinity (27). These receptors activate the cAMP/PKA signaling pathway and are positively coupled to G_αs_ (32). Hippocampal VPAC1 receptors can also couple to G_i/o_ proteins (33). VPAC2 receptors are primarily responsible for mediating VIP-induced enhancement of PCs excitability and NMDA receptor currents, which mainly occur post-synaptically and independently of GABAergic transmission (34). In human hippocampi surgically removed from patients with medically intractable TLE, a significant upregulation of VIP receptors on PCs has been observed (20), indicating an increased excitation of these cells by VIP. *In vitro* studies have shown that the selective VPAC1 antagonist PG 97-269 contributes to the inhibition of LTP following interictal-like activity in hippocampal slices (35). Additionally, a recent study in a lithium-pilocarpine mouse model demonstrated a compensatory decrease in VPAC1 receptor levels and an increase in VPAC2 receptor levels in hippocampal membranes (36), underscoring the critical role of VIP and its receptors in the pathophysiology of TLE. These reports align with our findings, which demonstrate both the protective effect of reduced VIP-INs activity and the intensified impact of heightened VIP-INs activity on epileptogenic markers in mice.

Several limitations need to be considered in relation to this study that. The primary limitation was the high variation in the number of seizures observed among animals, particularly in female mice, in the chronic model of epilepsy. However, this variation was expected (37). To address this issue and ensure the robustness of our findings, we also reported the number (and duration) of seizures as a ratio for each animal, comparing them to the number (and duration) of seizures before injection. As anticipated, individual female mice exhibited a greater degree of variation due to significant changes in hippocampal excitability during the estrous cycle (38, 39). Therefore, it is necessary to conduct additional research that specifically focuses on gender, taking into account the timing of KA injection in accordance with the estrous cycle of female mice.

Another limitation of the study concerns the sensitivity of targeting VIP-INs of the ventral subiculum through the injection of AAV-vector expressing TeLC. While our injection of AAV-TeLC was highly specific (over 90%), we were unable to target all VIP-INs in the subiculum, as reflected by a mean sensitivity of approximately 60%. Additionally, we observed variable expression of TeLC in other brain regions, including the ventral CA1, and occasionally in parts of the entorhinal cortex. Future studies utilizing alternative technologies with higher spatial precision may allow for more accurate targeting of the subiculum and its interneurons, including VIP-INs.

In our study, we employed an experimental model of acute focal seizures and a chronic model of epilepsy, both induced by IH-KA injection, to assess the role of VIP-INs of the ventral subiculum in the context of epilepsy. It is important to emphasize that conducting similar investigations in diverse animal models of epilepsy can provide additional valuable insights into the subject matter.

In conclusion, our study presents compelling evidence for the direct involvement of subicular VIP-INs in the pathogenesis of TLE. This finding represents a significant contribution to the field as it sheds light on the crucial role played by VIP-INs in subicular microcircuits and their potential as targets for therapeutic interventions aimed at enhancing seizure control. Future research employing techniques such as cell type-specific electrophysiology and *in vivo* imaging will further advance our understanding of subicular VIP-INs in the context of epilepsy. Moreover, these investigations will pave the way for the development of gene-targeted and cell type-specific therapies, offering promising avenues for more effective treatment strategies.

## Materials and Methods

### Mice

The experiments involving animals were conducted in strict adherence to national regulations and European Community laws. Approval for the animal experiments was obtained from the Committee for Animal Protection of the Austrian Ministry of Science. The VIP-cre transgenic mice (Vip^tm1(cre)Zjh^/J, RRID:IMSR_JAX:010908), which express Cre-recombinase under the VIP promoter, were initially obtained from The Jackson Laboratory through Charles River and were maintained on a C57BL/6N background. C57BL/6N wild-type mice were obtained from Charles River. The mice were housed in ventilated cages, with 3 to 5 mice per group, and had access to food and water *ad libitum.* Standard laboratory conditions were maintained, including a 12-hour light/dark cycle with lights on at 7:00 A.M. Male and female heterozygous mice aged 10 to 14 weeks were used for the experiments. For VIP co-localization studies only, VIP-cre homozygous mice were crossed with Ai9-tdTomato mice (B6.Cg-Gt(ROSA)^26Sortm9(CAG-tdTomato)Hze^/J; RRID:IMSR_JAX:007909) to generate VIP-cre:Ai9 double heterozygous mice.

### Preparation of the vectors

The AAV2/6 vectors carried either tetanus toxin light chain (TeLC) fused with a GFP tag or GFP alone. The reading frames for TeLC/GFP and GFP were inverted within a FLEX cassette allowing for AAV-mediated transgene expression specifically in VIP-INs at the injection site (40). Pseudotyped AAV2/6 vectors were produced and purified as previously described (41). In brief, HEK293 cells were co-transfected with rAAV plasmids pAM-FLEX-TeLC-GFP or pAM-FLEX-GFP, along with pDP1rs helper plasmids, using calcium phosphate co-precipitation. The resulting cleared cell lysates, treated with Benzonase, were purified by HPLC on AVB columns and subsequently subjected to dialysis. The AAV titers were determined by PCR as the number of genomic particles per milliliter (gp/ml), with titers of 1 × 10^12^ gp/ml for pAM-FLEX-TeLC-GFP and 1 × 10^13^ gp/ml for pAM-FLEX-GFP. To ensure the full activity of TeLC, we evaluated all seizure-related factors after 14 days following the injection, considering that the silencing effects would remain permanent thereafter (40).

For experiments involving DREADD (Designer Receptors Exclusively Activated by Designer Drugs) technology, we used AAV6-hSyn-DIO-hM3D(Gq)-mCherry (Addgene 44361), kindly provided by B. Roth and obtained from Addgene (USA). This vector was packaged into AAV2 capsids and injected with a final titer of 9 × 10^12^ gp/ml, as previously described in detail (42).

### Surgery

Stereotaxic surgeries were performed following previously described procedures (14). Male and female heterozygous VIP-cre transgenic mice aged 10–14 weeks were administered the analgesic drug meloxicam (5 mg/kg, s.c.) 15 min prior to surgery. Anesthesia was induced using 150 mg/kg of ketamine (50 mg/ml stock solution; Ketasol, Ogris Pharma Vertriebs-GmbH, Wels, Austria) via intraperitoneal injection, followed by maintenance anesthesia using 1–3% sevoflurane (3% for induction, 1–1.5% for maintenance; Sevorane, Abbott, IL, USA) administered through a veterinary anesthesia mask with oxygen (400 ml/min) as the carrier gas. The mice were secured in a stereotaxic frame (David Kopf Instruments, Tujunga, CA, USA), and the skin above the skull was opened.

A telemetry EEG transmitter (ETA-F10, Data Sciences International, St. Paul, MN, USA) was placed in a subcutaneous pocket on the left abdominal wall, with the electrode wires carefully pulled through a subcutaneous channel between the pocket and the exposed skull. The pocket was sutured after the procedure. Bilateral holes were drilled for AAV vector injection, insertion of an epidural recording electrode (coordinates from bregma: posterior 2.0 mm, lateral, 1.6 mm), and placement of an epidural reference electrode (coordinates from bregma: posterior 2.0 mm, lateral −1.6 mm). In a subset of animals, local field potential (LFP) was recorded to detect focal epileptiform activity. For these animals, a depth electrode (coordinates from bregma: posterior 1.9 mm, lateral 1.6 mm, ventral 1.9 mm) was inserted above the ipsilateral CA1 instead of the epidural recording electrode. The electrodes were secured to the skull using dental cement (Paladur, Heraeus Kulzer supplied by Henry Schein, Innsbruck, Austria).

Subsequently, 1 µL of AAV-TeLC-GFP or AAV-GFP was injected unilaterally into the transition zone of the left ventral subiculum and sector CA1 at a rate of 0.1 µl/min (coordinates from bregma: posterior 3.6 mm, lateral 3.2 mm, ventral 3.6 mm). The cannula remained in place for 10 minutes before slowly being retracted. In mice that underwent depth electrode recording, bilateral injections of AAV-TeLC-GFP or AAV-GFP were performed with the same coordinates as the unilateral injection (posterior 3.6 mm, lateral ± 3.2 mm, ventral 3.6 mm) to enhance the silencing effect of subicular VIP-INs in the subiculum. The AAV-TeLC-GFP or AAV-GFP injections were performed on the same days in mice from the same litters. Following surgery, the mice were housed individually.

For specific and reversible chemogenetic activation of VIP-INs in the subiculum, rAAV6-hSyn-DIO-hM3D(Gq) (0.5 µL) was injected at a rate of 0.1 µL/min into the left ventral subiculum of male and female heterozygous VIP-cre mice. The injections followed the same coordinates and protocol as the AAV-TeLC injections.

### Induction of the chronic model of epilepsy by IH-KA injection

To induce the chronic model of epilepsy, intrahippocampal injections of kainic acid (KA, Ascent Scientific, North Somerset, UK) were performed during transmitter implantation surgery in male and female heterozygous VIP-cre mice. KA (200 ng in 140 nL phosphate-buffered saline [PBS], pH 7.4) was injected at a speed of 140 nL/min using a 1 μL Hamilton syringe connected to a precision syringe pump (Nexus 3000 by Chemyx, Science Products, Hofheim, Germany). The injection cannula, a polyimide-coated silica capillary tubing with a 150 μm outer diameter (TSP075150, Polymicro Technologies, Phoenix, AZ, USA), was attached to the syringe using fine-bore polythene tubing (800/100/100; Smiths Medical International, Kent, UK). Following the injection, the cannula remained in place for 3 minutes before being slowly retracted. EEG recording was initiated immediately as the mice were transferred back to their cages. All mice in the study experienced acute seizures after the KA injection. The EEG activity returned to baseline after approximately 24 hours, indicating the end of status epilepticus (SE). As an exclusion criterion, mice that did not develop SRSs before vector injection (3 mice) were excluded from the rest of study.

### Experimental model of acute focal seizures induced by injection of IH-KA

To establish an experimental model of acute focal seizures, male and female heterozygous VIP-cre mice were anesthetized briefly with 3% sevoflurane, ten days after the injection of AAV-hM3Dq vector. Subsequently, they received an intrahippocampal injection of KA (7 ng in 500 nL phosphate-buffered saline [PBS], pH 7.4) at a speed of 500 nL/min using a 1 μL Hamilton syringe connected to a precision syringe pump. The injection cannula, a polyimide-coated silica capillary tubing with a 150 μm outer diameter, was attached to the syringe via fine-bore polythene tubing. After the injection, the cannula was left in place for 1 minute before being slowly retracted, and the mice were promptly transferred back to their cages for EEG recording.

Under these experimental conditions, KA induced acute seizures in 100% of the mice and the EEG activity returned to baseline after 1-2 hours.

### Transient activation of VIP-INs using the DREADD system

Seven days following the injection of AAV-hM3Dq, male and female heterozygous VIP-cre mice underwent the insertion of an EEG-transmitter, allowing for continuous telemetric EEG recording until the end of the study. Three days later (day 10 after vector injection), the selective hM3Dq agonist clozapine-N-oxide (CNO, Tocris Bioscience, Abingdon, UK) was intraperitoneally injected at a dose of 5 mg/kg. CNO was dissolved in sterile 0.9% NaCl at a concentration of 1.0 mg/ml. The control group of mice that received the AAV-hM3Dq vector injection were injected with saline instead of CNO.

Thirty minutes after CNO administration, acute focal-onset seizures were induced by intrahippocampal injection of KA. The effects of activating VIP-INs on the number and duration of seizures were assessed through telemetric EEG recordings.

### EEG and LFP recordings

Continuous EEG and LFP recordings were conducted using a telemetry system and Ponemah analysis software version 6.0.x. (Data Sciences International, St. Paul, MN, USA). The recordings were obtained at a sampling rate of 1000 Hz and saved onto external hard disk drives without any predetermined filter cutoff. To complement the analysis of behavior, a subset of animals was subjected to continuous video recordings. These recordings were captured using Axis 221 network cameras (Axis Communications, Lund, Sweden) along with infrared illumination (Conrad Electronics, Leonding, Austria).

### Analysis of epileptiform activity

For the analysis of epileptiform activity, two independent observers visually examined EEG and LFP traces using Ponemah analysis software version 6.0.x. (Data Sciences International, St. Paul, MN, USA). Seizures were identified as periods of uninterrupted activity with a minimum duration of 10 seconds and an amplitude at least twice that of the baseline, accompanied by postictal depression (EEG signal below baseline amplitude). In our cohort database, this signal corresponded to seizure Racine scale 3-5 (14), which was determined based on simultaneous video recordings.

To differentiate epileptiform spikes from normal baseline spikes or those caused by electrical or mechanical artifacts, specific features of each spike were analyzed, including amplitude, duration, frequency, and inter-spike intervals. An offline spike detection algorithm was implemented using a custom-made program developed in MATLAB 2020b (Mathworks, Natick, MA, USA). This program automatically detected spikes and spike trains based on threshold crossings. Initially, movement artifacts were automatically removed, and then a blind reviewer adjusted the spike detection threshold to account for variations in individual signal amplitude, ranging from 4 to 6 standard deviations. Epileptiform interictal spikes were defined as short (10 to 80 ms) high-amplitude discharges above the threshold, with a minimum peak distance of 150 ms. Detected spikes were visually confirmed by the reviewers.

Spike trains were defined as sequences of three or more spikes occurring with a frequency greater than 1 Hz and lasting for at least 1 second (43). Since postictal depression is not always visible during SE, in addition to comparing the groups based on the number of seizures, the severity of SE was also assessed by analyzing the number of epileptiform spikes and spike trains (44).

### Wake/Sleep annotation

We employed our previously published algorithm to automatically differentiate between sleep and wakefulness in single-channel EEG recordings by utilizing high-frequency oscillations (> 200 Hz). This approach achieved an accuracy of over 90 percent (45). Subsequently, we employed the SCOPRISM algorithm (46) to differentiate between rapid eye movement sleep (REMS) and non-rapid eye movement sleep (NREMS). The original SCOPRISM algorithm suggests dividing theta by delta and employing a threshold of 0.75, classifying values above the threshold as REMS and those below as NREMS. However, in our cohort database, where the vigilance state is manually determined based on EEG analysis from the visual, somatosensory, and motor cortex, as well as the EMG signal, we observed that a threshold of 0.8 produced more accurate results. Consequently, we decided to utilize this threshold for the remainder of the study.

For epileptic animals, we compared the vigilance state not only after vector injection (during a 24-hour period 14 days after vector injection), but also before vector injection to establish a baseline for the groups’ vigilance patterns. To assess the general sleep architecture, we derived the proportion of vigilance states for non-overlapping two-hour observation episodes throughout the 24-hour period, as well as the distribution of bout lengths for each vigilance state, from the scoring vectors.

### Double- and triple-labeling by immunofluorescence

At the end of the experiment, which typically occurred 4-8 weeks after the initial vector injection, the mice were deeply anesthetized and euthanized with thiopental (150 mg/kg). Perfusion with 4% paraformaldehyde was then performed to fix the brains. Horizontal sections with a thickness of 40 μm were prepared for the immunofluorescence study.

Monoclonal rat anti-GFP (1:2000; catalog #04404-84, Nacalai Tesque, obtained through GERBU Biotechnik; RRID:AB_10013361) and rabbit anti-VIP (1:500; catalog #20077, Immunostar, obtained through ACRIS, RRID: AB_572270) were used as primary antibodies. The antibodies had been previously characterized by the supplier or validated in previous experiments (16). For the double labeling of GFP and VIP, the horizontal sections were incubated with 10% normal goat or horse serum (catalog #26050088, Gibco, through Thermo Fischer) in Tris-HCl-buffered saline (TBS; 50 mm), pH 7.2, containing 0.4% Triton X-100 (TBS-Triton) for 90 minutes. Subsequently, the sections were incubated with the respective primary antisera for 16 hours at room temperature.

After washing with TBS-Triton, a secondary reaction was performed by simultaneously incubating the sections with a donkey antibody directly conjugated to Alexa Fluor 488 (1:500; catalog #A-21208, Molecular Probes) for GFP and a biotin-coupled goat anti-rabbit antibody (1:500; catalog #A-16027, Invitrogen) for VIP at room temperature for 120 minutes. The sections were then washed with TBS and incubated in streptavidin (1:100; catalog #SA 5649, Vektor) for 100 minutes. Following another wash with TBS, the sections were mounted on slides in gelatin and covered with mounting medium (86% glycerol and 2.5% DABCO in PBS).

To characterize the three subgroups of subicular VIP-INs, we utilized VIP:Ai9-tdTomato mice. These mice were deeply anesthetized with thiopental (150 mg/kg) and perfused to prepare 40-μm-thick coronal sections for immunofluorescence analysis. We already reported a high level of colocalization between tdTomato and VIP in VIP:Ai9-tdTomato mice, indicating a sensitivity and specificity exceeding 90% (47). For the immunofluorescence labeling of VIP (tdTomato), Calretinin (CR), and Cholecystokinin (CCK), the free-floating coronal sections were incubated with 10% normal goat serum (catalog #5425, Cell Signaling) in Tris-HCl-buffered saline, pH 7.2, containing 0.4% Triton X-100 (TBS-Triton) for 90 minutes at room temperature. Subsequently, the sections were incubated with the respective primary antisera for 20 h at 4°C. The primary antibodies used were rabbit anti-CR antibody (1:1000; catalog 7769/3H, SWant) and guinea pig anti-CCK antibody (1:1000; catalog #438-004; Synaptic Systems).

After washing with TBS-Triton, a secondary reaction was performed by simultaneously incubating the sections with a rabbit antibody directly conjugated to Alexa Fluor 488 (1:500; catalog #4412S, Cell Signaling) for CR and with a guinea pig antibody directly conjugated to Alexa Fluor 647 (1:500; catalog #ab150187, Abcam) for CCK at room temperature for 120 minutes. The sections were then washed with TBS and mounted on slides with gelatin. Finally, the slides were covered with mounting medium composed of 86% glycerol and 2.5% DABCO in PBS.

To determine the number of neurons co-expressing TeLC/GFP and VIP at the AAV vector injection site, microphotographs of 40-μm-thick horizontal double immunofluorescence-labeled sections (four sections per mouse, ten mice) were captured at 20× primary magnification using a fluorescence microscope (Imager.M1, Carl Zeiss). The acquired images were imported into NIH ImageJ 1.51d (National Institutes of Health, Bethesda, MD, USA), and the photographs of the individual channels were displayed side by side for analysis. The specific area of interest for evaluation was defined by the region containing GFP-positive neurons, which were directly affected by the vector injection. The numbers of single-labeled (TeLC/GFP or VIP) and double-labeled (co-expressing TeLV/GFP and VIP) cells were manually determined by visual inspection of the images.

### Statistical Analyses

Statistical analyses were performed using GraphPad Prism statistical software (version 9.1.1; GraphPad Software). Due to the significant variation in seizure numbers observed in the IH-KA model (37), the ratio of seizures for each animal was calculated in addition to reporting the raw number and duration of seizures. The ratio was determined by comparing the number (or duration) of seizures after vector injection to the number (or duration) of seizures before injection using the formula (seizures after - seizures before) divided by (seizures after + seizures before).

The differences between the two groups were analyzed using the non-parametric Mann–Whitney U test. To compare seizures, epileptiform spikes, spike trains, and sleep-related factors before and after vector injection over a specific time period, a two-way repeated-measures ANOVA was conducted, followed by Šídák’s multiple-comparison post hoc test. A p-value of less than 0.05 was considered statistically significant. Graphs were generated using GraphPad Prism 9.1.1 and MATLAB 2020b.

## Acknowledgments

Valuable insight into the hippocampal disinhibitory circuit and its possible relevance to the findings of the current experiment was provided by Professor Lisa Topolnik from Laval University, Canada. She is deserving of our sincere thanks. We would like to express our sincere gratitude to Professor Francesco Ferraguti, Medical University of Innsbruck (MUI), Austria, for generously providing three VIP:Ai9-tdTomato mice for the experiment. We also want to thank Lisa Bergmeister (MUI), Pradeepa Mohan (MUI) and Dr. Samira Zahmatkesh (University of Innsbruck, Austria) for their fruitful discussions and knowledgeable suggestions regarding the statistics and representation of the results. The authors thanks Atea Shkodra for her excellent technical assistance. This research was funded in whole, or in part, by the Austrian Science Fund (FWF) [P 31777 to M.D.]. For the purpose of open access, the author has applied a CC BY public copyright licence to any Author Accepted Manuscript version arising from this submission.

## Data availability

The EEG recordings and all algorithms implemented in this work can be made available upon request by contacting the first author (Dr. Sadegh Rahimi, Medical University of Innsbruck, Austria).

## Author contributions

S.R. Data curation; Formal analysis; Investigation; Methodology; Visualization; Software; Writing - original draft; Writing - review & editing.

P.S. Formal analysis; Software; Writing - review & editing.

P.M. Data curation.

A.S. Formal analysis; Validation; Data curation.

A.R.-P. Visualization; Writing - review & editing.

R.T. Resources.

M.D. Conceptualization; Investigation; Methodology; Project administration; Supervision; Validation; Writing - review & editing.

## Declaration of generative AI and AI-assisted technologies in the writing process

During the preparation of this work, Dr. Sadegh Rahimi, first author of the manuscript, used *Wordtune* and *Chat Generative Pre-trained Transformer (ChatGPT)* in order to improve the language of the manuscript. After using these services, Dr. Meinrad Drexel and Dr. Sadegh Rahimi reviewed and edited the content as needed and takes full responsibility for the content of the publication.

---

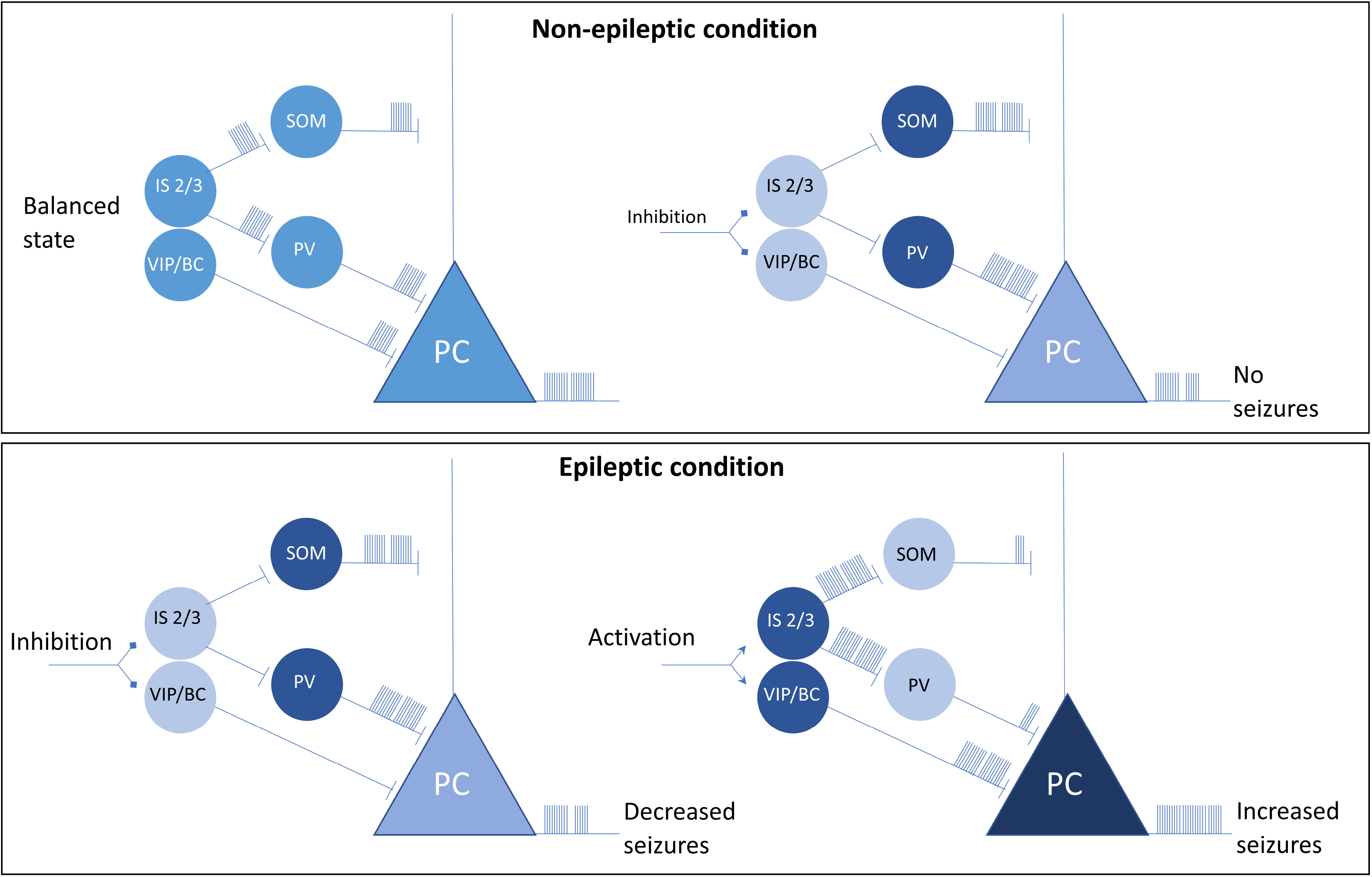

**Supplementary Figure 1.**
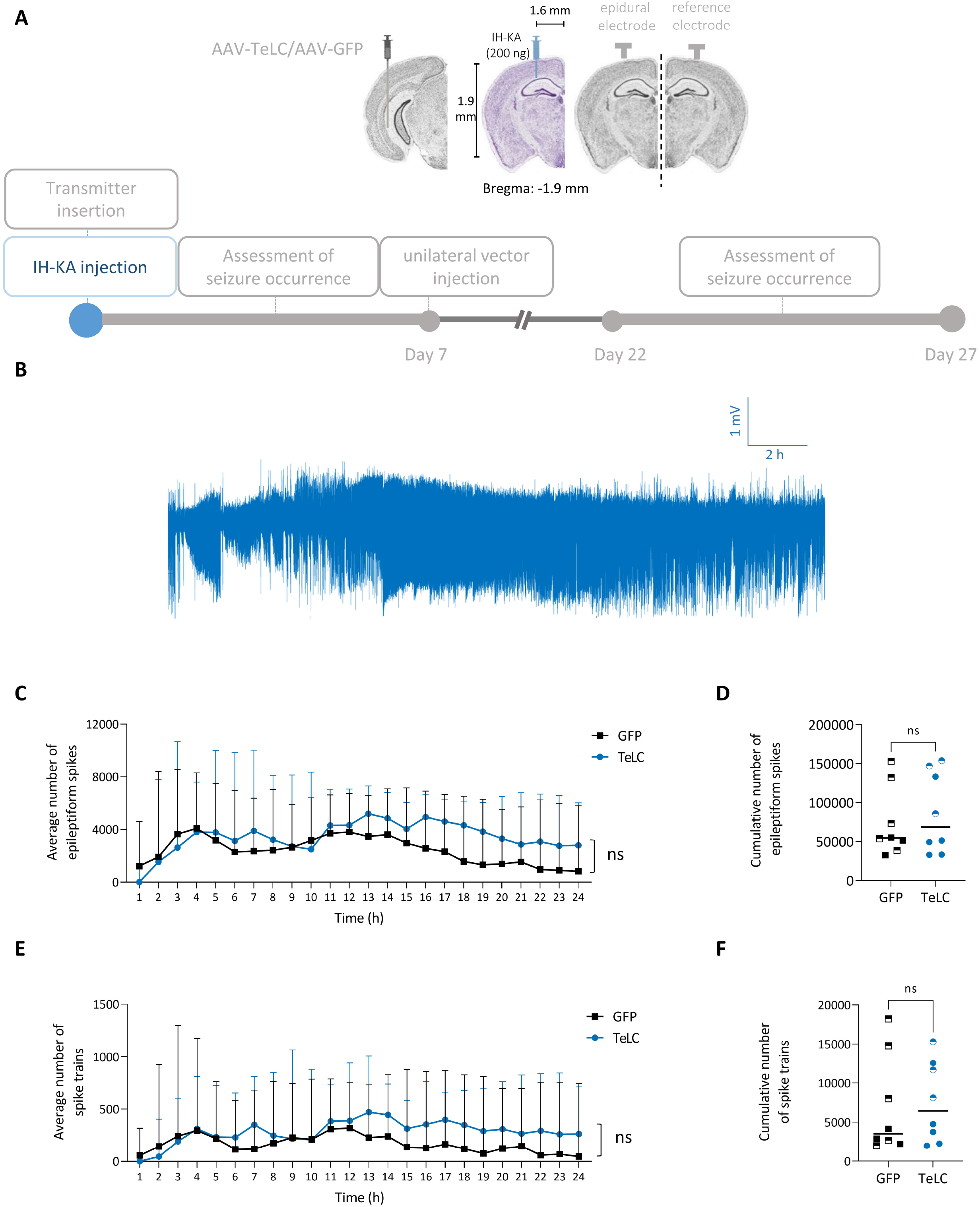
No difference was observed in severity of SE between groups. (A) The study protocol and the coordinates and for intrahippocampal injection of KA (IH-KA) (*n*=16). The animals were blindly divided into two groups, GFP (*n*=8) and TeLC (*n*=8), receiving AAV-GFP or AAV-TeLC after 7 days, respectively. (B) Representative example of status epilepticus (SE) in the first 24 hours after intrahippocampal KA injection. Postictal depression is not always visible during SE. (C) During SE, the median (range) hourly number of epileptiform spikes was 3149 (1786 to 4071) for the GFP group (n=8) and 3767 (206.8 to 4633) for the TeLC group (*n*=8). No significant difference was found between the groups according to a two-way ANOVA test (*F* = 0.252, *p* = 0.623). (D) The median (range) number of epileptiform spikes in the GFP group (*n*=8) was 54511 (32651 to 153106), and for the TeLC group (n=8) it was 68674 (33018 to 153892). There was no significant difference between the groups according to a two-tailed Mann-Whitney test (*p* = 0.878). (E) During SE, the median (range) hourly number of spike trains was 283 (110.4 to 401.6) for the GFP group (N=8) and 324.2 (6.62 to 471.90) for the TeLC group (*n*=8). No significant difference was found between the groups according to a two-way ANOVA test (*F* = 0.058, *p* = 0.813). (F) The median (range) number of spike trains was 3519 (1993 to 18226) for the GFP group (*n*=8) and 6439 (1996 to 15306) for the TeLC group (*n*=8). There was no significant difference between the groups according to a two-tailed Mann-Whitney test (*p* = 0.645). Male individuals are represented by half-filled shapes in both groups. Mouse coronal slices in (A) are adapted from the Allen Mouse Brain Atlas, available online: http://atlas.brain-map.org/atlas

**Supplementary Figure 2.**
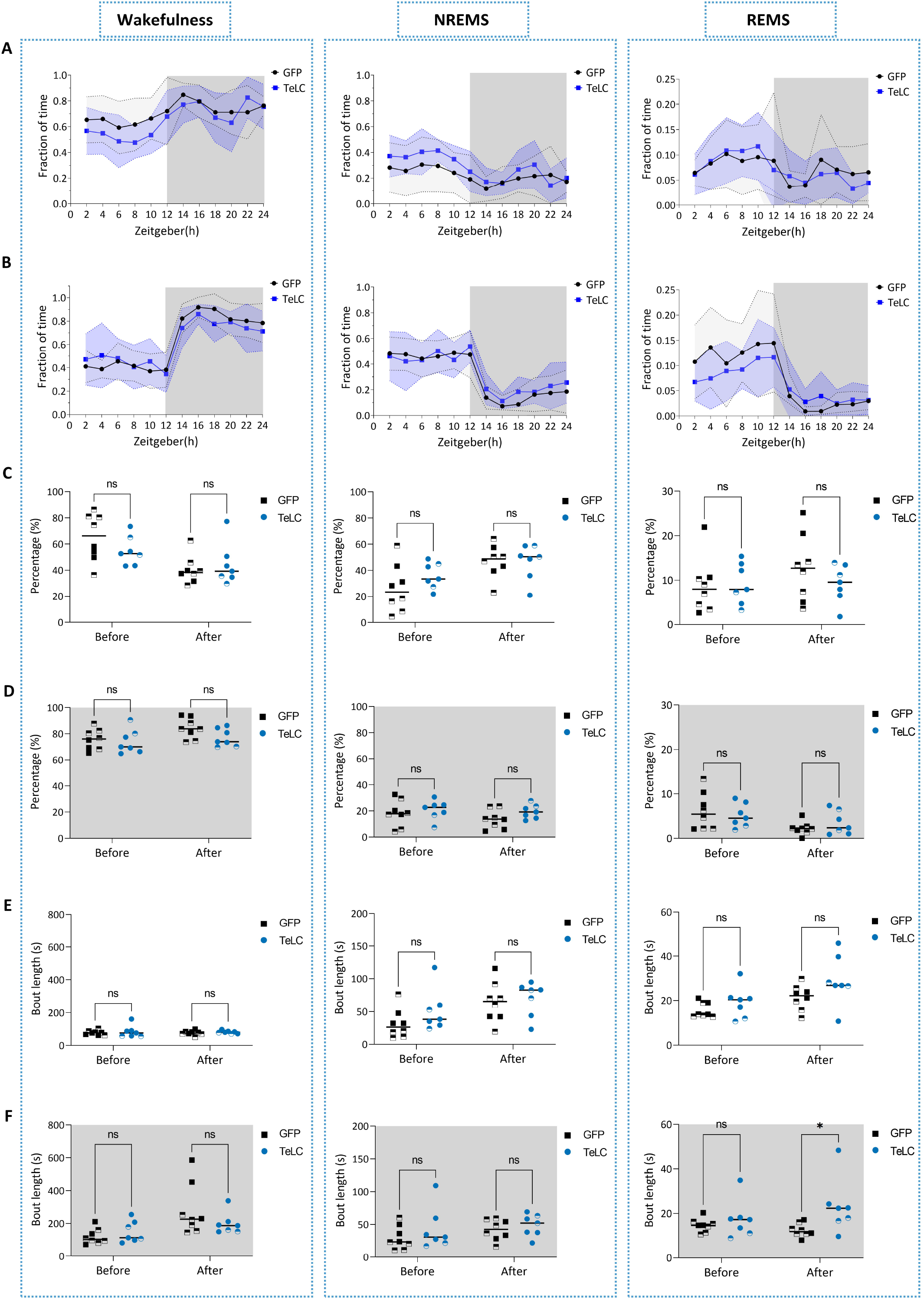
Silencing of VIP-INs of the ventral subiculum in epileptic mice causes no changes in sleep pattern. (A) Before vector injection (three days after intrahippocampal KA injection [IH-KA]), both groups exhibited altered patterns in the three vigilance states due to epilepsy, resulting in the disappearance of the typical nocturnal pattern of wakefulness. (B) Two weeks after vector injection (Day 22), both groups showed a restoration of the nocturnal pattern of wakefulness and a recovery in sleep architecture. (C) During the light phase, before vector injection, there were no significant differences in the percentage of wakefulness, non-rapid eye movement sleep (NREMS), and rapid eye movement sleep (REMS) between the GFP (*n*=8) and TeLC (*n*=7, since signal from one mouse was not fit to our sleep scoring algorithm) groups. Also after vector injection, there were no significant differences observed in the percentage of wakefulness, NREMS, and REMS between the groups. Data were analyzed by fitting a mixed model. (D) During the dark phase, before and after vector injection, there were no significant differences in the percentage of wakefulness, NREMS, and REMS between the GFP (*n*=8) and TeLC (*n*=7, since signal from one mouse was not fit to our sleep scoring algorithm) groups. Data were analyzed by fitting a mixed model. (E) During the light phase, before vector injection, there were no significant differences in the bout length of wakefulness, NREMS, and REMS between the GFP (*n*=8) and TeLC (*n*=7, since signal from one mouse was not fit to our sleep scoring algorithm) groups. Also after vector injection, there were no significant differences observed in bout length of wakefulness, NREMS, and REMS between the groups. Data were analyzed by fitting a mixed model. (F) During the dark phase, before vector injection, there were no significant differences in the bout length of wakefulness, NREMS, and REMS between the GFP (*n*=8) and TeLC (*n*=7) groups. After vector injection, there were no significant differences observed in the bout length of wakefulness and NREMS between the two groups, but a significant difference was observed in the bout length of REMS (*p* = 0.02, Šídák post hoc test). However, analyzed by fitting a mixed model, also for REMS, no difference was observed between groups (F = 2.77, *p =* 0.12). Male individuals are represented by half-filled shapes in both groups.

## References

1. Fei F, Wang X, Wang Y, Chen Z. Dissecting the role of subiculum in epilepsy: Research update and translational potential. Prog Neurobiol 2021; 201:102029.

2. Cohen I, Navarro V, Clemenceau S, Baulac M, Miles R. On the origin of interictal activity in human temporal lobe epilepsy in vitro. Science 2002; 298(5597):1418–21.

3. Wozny C, Knopp A, Lehmann T-N, Heinemann U, Behr J. The subiculum: a potential site of ictogenesis in human temporal lobe epilepsy. Epilepsia 2005; 46 Suppl 5:17–21.

4. Blümcke I, Thom M, Aronica E, Armstrong DD, Bartolomei F, Bernasconi A et al. International consensus classification of hippocampal sclerosis in temporal lobe epilepsy: a Task Force report from the ILAE Commission on Diagnostic Methods. Epilepsia 2013; 54(7):1315–29.

5. Drexel M, Preidt AP, Sperk G. Sequel of spontaneous seizures after kainic acid-induced status epilepticus and associated neuropathological changes in the subiculum and entorhinal cortex. Neuropharmacology 2012; 63(5):806–17. Available from: URL: https://www.sciencedirect.com/science/article/pii/s0028390812002675.

6. Chung S, Spruston N, Koh S. Age-dependent changes in intrinsic neuronal excitability in subiculum after status epilepticus. PLoS ONE 2015; 10(3):e0119411.

7. Fei F, Wang X, Xu C, Shi J, Gong Y, Cheng H et al. Discrete subicular circuits control generalization of hippocampal seizures. Nat Commun 2022; 13(1):5010.

8. Lippmann K, Klaft Z-J, Salar S, Hollnagel J-O, Valero M, Maslarova A. Status epilepticus induces chronic silencing of burster and dominance of regular firing neurons during sharp wave-ripples in the mouse subiculum. Neurobiol Dis 2022; 175:105929.

9. Bezaire MJ, Soltesz I. Quantitative assessment of CA1 local circuits: knowledge base for interneuron-pyramidal cell connectivity. Hippocampus 2013; 23(9):751–85.

10. Kenneth A. Pelkey, Ramesh Chittajallu, […], and Chris J. McBain. Hippocampal GABAergic Inhibitory Interneurons.

11. Kepecs A, Fishell G. Interneuron cell types are fit to function. Nature 2014; 505(7483):318–26.

12. Knopp A, Frahm C, Fidzinski P, Witte OW, Behr J. Loss of GABAergic neurons in the subiculum and its functional implications in temporal lobe epilepsy. Brain 2008; 131(Pt 6):1516–27.

13. Drexel M, Preidt AP, Kirchmair E, Sperk G. Parvalbumin interneurons and calretinin fibers arising from the thalamic nucleus reuniens degenerate in the subiculum after kainic acid-induced seizures. Neuroscience 2011; 189(1-2):316–29.

14. Drexel M, Romanov RA, Wood J, Weger S, Heilbronn R, Wulff P et al. Selective Silencing of Hippocampal Parvalbumin Interneurons Induces Development of Recurrent Spontaneous Limbic Seizures in Mice. J Neurosci 2017; 37(34):8166–79.

15. Dinocourt C, Petanjek Z, Freund TF, Ben-Ari Y, Esclapez M. Loss of interneurons innervating pyramidal cell dendrites and axon initial segments in the CA1 region of the hippocampus following pilocarpine-induced seizures. J Comp Neurol 2003; 459(4):407–25.

16. Drexel M, Rahimi S, Sperk G. Silencing of Hippocampal Somatostatin Interneurons Induces Recurrent Spontaneous Limbic Seizures in Mice. Neuroscience 2022; 487:155–65.

17. Klausberger T, Somogyi P. Neuronal diversity and temporal dynamics: the unity of hippocampal circuit operations. Science 2008; 321(5885):53–7.

18. Pelkey KA, Chittajallu R, Craig MT, Tricoire L, Wester JC, McBain CJ. Hippocampal GABAergic Inhibitory Interneurons. Physiol Rev 2017; 97(4):1619–747.

19. Kullander K, Topolnik L. Cortical disinhibitory circuits: cell types, connectivity and function. Trends Neurosci 2021; 44(8):643–57.

20. Lanerolle NC de, Gunel M, Sundaresan S, Shen MY, Brines ML, Spencer DD. Vasoactive intestinal polypeptide and its receptor changes in human temporal lobe epilepsy. Brain research 1995; 686(2):182–93.

21. Ko FJ, Chiang CH, Liu WJ, Chiang W. Somatostatin, substance P, prolactin and vasoactive intestinal peptide levels in serum and cerebrospinal fluid of children with seizure disorders. Gaoxiong Yi Xue Ke Xue Za Zhi 1991; 7(8):391–7.

22. Liang C, Zhang C-Q, Chen X, Wang L-K, Yue J, An N et al. Differential Expression Hallmarks of Interneurons in Different Types of Focal Cortical Dysplasia. J Mol Neurosci 2020; 70(5):796–805. Available from: URL: https://link.springer.com/article/10.1007/s12031-020-01492-0.

23. Hall S, Hunt M, Simon A, Cunnington LG, Carracedo LM, Schofield IS et al. Unbalanced Peptidergic Inhibition in Superficial Neocortex Underlies Spike and Wave Seizure Activity. J Neurosci 2015; 35(25):9302–14.

24. Henkel ND, Smail MA, Wu X, Enright HA, Fischer NO, Eby HM et al. Cellular, molecular, and therapeutic characterization of pilocarpine-induced temporal lobe epilepsy. Sci Rep 2021; 11(1):19102.

25. Wyeth M, Buckmaster PS. Lack of Hyperinhibition of Oriens Lacunosum-Moleculare Cells by Vasoactive Intestinal Peptide-Expressing Cells in a Model of Temporal Lobe Epilepsy. eNeuro 2021; 8(6).

26. David LS, Topolnik L. Target-specific alterations in the VIP inhibitory drive to hippocampal GABAergic cells after status epilepticus. Exp Neurol 2017; 292:102–12.

27. Cunha-Reis D, Caulino-Rocha A. VIP Modulation of Hippocampal Synaptic Plasticity: A Role for VIP Receptors as Therapeutic Targets in Cognitive Decline and Mesial Temporal Lobe Epilepsy. Front Cell Neurosci 2020; 14:153.

28. Moshé SL, Perucca E, Ryvlin P, Tomson T. Epilepsy: new advances. Lancet 2015; 385(9971):884–98.

29. Rowley NM, Madsen KK, Schousboe A, Steve White H. Glutamate and GABA synthesis, release, transport and metabolism as targets for seizure control. Neurochem Int 2012; 61(4):546–58. Available from: URL: https://www.sciencedirect.com/science/article/pii/s0197018612000642.

30. Kang Y-J, Clement EM, Park I-H, Greenfield LJ, Smith BN, Lee S-H. Vulnerability of cholecystokinin-expressing GABAergic interneurons in the unilateral intrahippocampal kainate mouse model of temporal lobe epilepsy. Exp Neurol 2021; 342:113724.

31. Acsády L, Arabadzisz D, Freund TF. Correlated morphological and neurochemical features identify different subsets of vasoactive intestinal polypeptide-immunoreactive interneurons in rat hippocampus. Neuroscience 1996; 73(2):299–315.

32. Yang K, Lei G, Jackson MF, Macdonald JF. The involvement of PACAP/VIP system in the synaptic transmission in the hippocampus. J Mol Neurosci 2010; 42(3):319–26.

33. Martin Shreeve S. Identification of G-proteins coupling to the vasoactive intestinal peptide receptor VPAC(1) using immunoaffinity chromatography: evidence for precoupling. Biochem Biophys Res Commun 2002; 290(4):1300–7.

34. Cunha-Reis D, Sebastião AM, Wirkner K, Illes P, Ribeiro JA. VIP enhances both pre- and postsynaptic GABAergic transmission to hippocampal interneurones leading to increased excitatory synaptic transmission to CA1 pyramidal cells. Br J Pharmacol 2004; 143(6):733–44.

35. Carvalho-Rosa JD, Cunha-Reis D. Endogenous VIP VPAC1 receptor activation during ictal and interictal-like activity induced in vitro by biccuculine and 0-Mg2+ modulates subsequent LTP expression in the rat hippocampus. Front Cell Neurosci 2019; 13.

36. Serpa A, Bento M, Caulino-Rocha A, Pawlak S, Cunha-Reis D. Opposing reduced VPAC1 and enhanced VPAC2 VIP receptors in the hippocampus of the Li2+-pilocarpine rat model of temporal lobe epilepsy. Neurochem Int 2022; 158:105383.

37. Rusina E, Bernard C, Williamson A. The Kainic Acid Models of Temporal Lobe Epilepsy. eNeuro 2021; 8(2).

38. Scharfman HE, Mercurio TC, Goodman JH, Wilson MA, MacLusky NJ. Hippocampal excitability increases during the estrous cycle in the rat: a potential role for brain-derived neurotrophic factor. J. Neurosci. 2003; 23(37):11641–52.

39. Li J, Leverton LK, Naganatanahalli LM, Christian-Hinman CA. Seizure burden fluctuates with the female reproductive cycle in a mouse model of chronic temporal lobe epilepsy. Exp Neurol 2020; 334:113492.

40. Murray AJ, Sauer J-F, Riedel G, McClure C, Ansel L, Cheyne L et al. Parvalbumin-positive CA1 interneurons are required for spatial working but not for reference memory. Nat Neurosci 2011; 14(3):297–9.

41. Mietzsch M, Broecker F, Reinhardt A, Seeberger PH, Heilbronn R. Differential adeno-associated virus serotype-specific interaction patterns with synthetic heparins and other glycans. J Virol 2014; 88(5):2991–3003.

42. Tasan RO, Nguyen NK, Weger S, Sartori SB, Singewald N, Heilbronn R et al. The central and basolateral amygdala are critical sites of neuropeptide Y/Y2 receptor-mediated regulation of anxiety and depression. J Neurosci 2010; 30(18):6282–90.

43. Auer T, Schreppel P, Erker T, Schwarzer C. Functional characterization of novel bumetanide derivatives for epilepsy treatment. Neuropharmacology 2020; 162:107754.

44. Sharma S, Puttachary S, Thippeswamy A, Kanthasamy AG, Thippeswamy T. Status Epilepticus: Behavioral and Electroencephalography Seizure Correlates in Kainate Experimental Models. Front. Neurol. 2018; 9.

45. Rahimi S, Soleymankhani A, Joyce L, Matulewicz P, Kreuzer M, Fenzl T et al. Discriminating rapid eye movement sleep from wakefulness by analyzing high frequencies from single-channel EEG recordings in mice. Sci Rep 2023; 13(1):9608.

46. Bastianini S, Berteotti C, Gabrielli A, Del Vecchio F, Amici R, Alexandre C et al. SCOPRISM: a new algorithm for automatic sleep scoring in mice. J Neurosci Methods 2014; 235:277–84.

47. Ramos-Prats A, Paradiso E, Castaldi F, Sadeghi M, Mir MY, Hörtnagl H et al. VIP-expressing interneurons in the anterior insular cortex contribute to sensory processing to regulate adaptive behavior. Cell Rep 2022; 39(9):110893.

